# Automated High-Throughput Microfluidic Platform for Cell Pairing and Chemical Fusion

**DOI:** 10.64898/2026.10.02.756271

**Authors:** Robinzon Silvestre-de-León, Everardo González-González, Roberto Rodriguez-Mocayo, José Antonio Cruz-Cardenas, Sergio O. Martinez-Chapa, Joel Voldman, Mohammad Mahdi Aeinehvand

## Abstract

Cell fusion enables cell-based applications, yet systematic optimization remains constrained by low throughput (microfluidic devices) and manual operation (bulk protocols). Here, an automated microfluidic platform is presented for cell pairing and chemical fusion at a statistically relevant scale. High throughput is achieved by arranging ~350,000 deformability-based cell traps as concentric circular arrays in a radial chip. Using tumor and dendritic cells as a model for cancer vaccine, the device reproducibly co-occupies ~75% of traps (~250,000) of which ~140,000 hold exactly one cell of each type. This exceeds the yield of previously reported in hydrodynamic pairing arrays by more than an order of magnitude. A concentration step-up-and-down PEG profile yielded a fusion efficiency of ~20% (range 17.7-23.4%, n =3) at viability above 90% that results in producing at least ~50,000 hybrids per chip. This efficiency is comparable to that reported for PEG-based fusion in small-scale microfluidic devices but obtained at higher throughput and approximately four-fold higher than the conventional benchtop protocol. Fusion efficiencies were quantified by image analysis and validated by flow cytometry. By combining automation, large-area circular trap arrays and reagent control, the platform is useful for data-rich optimization of cell engineering processes previously constrained by limited microfluidic throughput.

## 1. Introduction

Cell fusion is a fundamental process with broad relevance in biology and health sciences implicated in the creation of hybrid cells for antibody production, cell reprogramming studies, cancer research, and therapeutic discovery.^[1–5]^ In cancer immunotherapy, fusion of dendritic cells with tumor cells yields hybrids that present tumor antigen while retaining antigen-presenting function. ^[2,4,6]^ Despite its importance, the most common fusion methods such as chemical fusion and electrofusion, performed in chambers or tubes, remain limited in compatibility with automation and therefore scalability. Additionally, these methods do not allow to achieve organized cell pairing that is required for high fusion efficiency.^[7,8]^

Conventional chemical fusion protocols based on polyethylene glycol (PEG) are widely used because of their simplicity, low cost, and high yield when abundant cells are available. However, bulk chemical fusion methods suffer from random cell–cell contact and heterogeneous exposure to PEG, which limits fusion efficiency and reproducibility.^[5,9–12]^ Attempts to mitigate this by physically controlling PEG delivery, for example transient-pressure jet injection, illustrate how strongly the outcome depends on exposure control.^[13]^ Electrofusion avoids PEG toxicity by using electrical pulses to create pores and reversibly permeabilize the cell membranes which can also raise the fusion efficiency.^[14,15]^ It nonetheless requires electrode patterning and alignment, a synchronized pulse generator and high local field strengths, which add instrumentation overhead in screening workflows and additional technical knowledge in clinical manufacturing pipelines.^[16,17]^

Recent advances in microfluidics have begun to address these limitations by enabling microscale confinement, precise reagent handling, and controlled positioning of individual cells for their pairing. ^[3,10,18–21]^ In addition, since the majority of microfluidic chips are fabricated with PDMS and glass slides, these technologies are compatible with optical microscopy for image analysis, which enables downstream experiments (e.g., dynamic monitoring) and facilitates automation.

Several microfluidic platforms have been reported for electrical and optical fusion,^[10,18,22,23]^ and for chemical fusion using controlled PEG exposure.^[3,24]^ Hydrodynamic devices that capture and pair cells in traps within straight channels provide reproducible control of the pairing step. This allows to reach fusion efficiencies of 40–60% with electrofusion and 3–30% using PEG-based protocols.^[3,18]^ Reported arrays of this type contain at most ~ 10,000 traps^[3,18,24]^ which bounds the number of pairs obtainable per chip. Scaling these architectures is constrained by the driving pressure drop required to sustain a given flow rate through a straight channel, which grows in direct proportion to the length as more traps are added in series.^[25]^ Moreover, in practice, sequential trap filling from inlet to outlet produces non-uniform loading that worsens as the array is extended.^[26]^ Dielectrophoretic microfluidic approaches that pair cells in traps, ^[22]^ wells^[23,27]^ or through orifices^[19,28]^ achieve the highest reported fusion efficiencies (up to 90%). Their integration requires patterned electrode arrays, an alignment-sensitive bonding step, and a synchronized function generator. In addition, the medium must be exchanged for a low-conductivity buffer before pulsing, so an automated sequence must accommodate additional wash and buffer-exchange stages. Droplet approaches^[20,21]^ generate hundreds of thousands of cell pairs in droplets and are attractive for their throughput. However, the surfactant-stabilized interface makes stepwise medium and PEG exchange difficult for chemical fusion protocols that depend on timed washout. ^[20,29]^ Electrofusion inside droplets circumvents the PEG exchange problem, yet reported efficiencies remain low, possibly because the contact point between partners is uncontrolled, and pulse delivery must be synchronized with the droplet generation frequency.^[21]^ Therefore, to our knowledge, no reported architecture combines cell pairing at the 10^5^ scale with the timed reagent exchange across a fully occupied array that a systematic test of graded PEG delivery requires.

Here we report an automated high-throughput cell pairing microfluidic platform for chemical cell fusion. It contains ~350,000 cell traps, which enables processing of hundreds of thousands of trapped cells on a single PDMS chip. The traps are arranged in a circular format within a radially symmetric chip. This layout was chosen for ease of future adaptation to centrifugal microfluidic platforms, and because a single central port feeds the entire array without a distribution manifold. Automation of the injection sequence removes the operator from every timed step from cell loading through cell fusion, with priming of the tubing and reservoirs carried out beforehand. The reagent exchange is executed with a reproducibility that manual processes, which vary among users, cannot achieve. We used this scale to test if concentration stepwise PEG delivery, rather than single-step exposure improves fusion yield while preserving viability. PEG cytotoxicity scales with both concentration and exposure time, and sequential PEG treatments are known to improve fusion efficiency relative to single-step exposure.^[30,31]^ Graded delivery should therefore reach fusogenic conditions while limiting the abruptness of the osmotic transition. On this basis, concentration step-based PEG protocols were evaluated as a strategy to optimize cell fusion and preserve viability. Finally, to establish utility for therapeutic applications, we fused immortalized mouse dendritic cells (MutuDC1940-GFP) with tumor cells (C1498-RFP) and demonstrate controlled cell trapping and pairing, efficient chemical fusion, and recovery of viable hybrid cells (up to 50,000 per chip). This work develops a platform and establishes a methodology for an automated and high-throughput single-cell trapping, pairing and optimization of cell engineering parameters that bridges microscale precision and the sample sizes required for statistically powered optimization.

## 2. Results

### 2.1. Chip architecture and radial flow distribution

The automated cell pairing platform uses a microfluidic chip with cell traps arranged in a concentric ring arrays oriented perpendicular to the central port (**Figure 1a**). The details for the array building and some scale-up notes are described in Supplementary Information S1 and S2. The circular layout solves the distribution problem that constrains wide linear arrays. A linear array wide enough to hold hundreds of thousands of traps would require a branched manifold to deliver flow uniformly across the channel width, which adds fabrication complexity and compromises footprint. In this device, liquid is injected through a single port at the center of the chip and leaves radially through a horizontal outlet that runs around the outer rim, so that flow is distributed symmetrically along every radius instead of along a single axis. Consequently, the flow velocity is uniform among all traps belonging to the same circular array, whereas a monotonic gradient persists across the radius. Per-trap flow and wall shear fall as 1/r, so they differ here by the ratio r_2_/r_1 ≈_ 13 between the innermost and outermost rings. At the flow rates used here, this gradient does not compromise operation, because deformability-based trapping tolerates a range of flow rates depending on both cell and trap properties and sizes. The functionality of the circular arrays with the inherent gradient of flow across chip radius was first tested and validated by a pair of syringe pumps and manual steps as briefly described in Supplementary Information S3. The highest local velocities occur on the innermost rings, and this margin narrows as the operating flow rate rises. Where a wider operating range is required, the gradient can be compensated by grading the constriction with radius or by using a thinner trap-zone ring of larger radius, both of which are described in a prior patent application on this architecture.^[32]^ The pressure cost of enlarging the array is also fundamentally different. Whereas the driving pressure of a straight channel grows in direct proportion to its length, the radial geometry accumulates pressure only logarithmically with radial span, ΔP ∝ ln(r_2_/r_1_). As a result, a tenfold increase in trap count raises ΔP by less than a factor of two.

**Figure 1.**
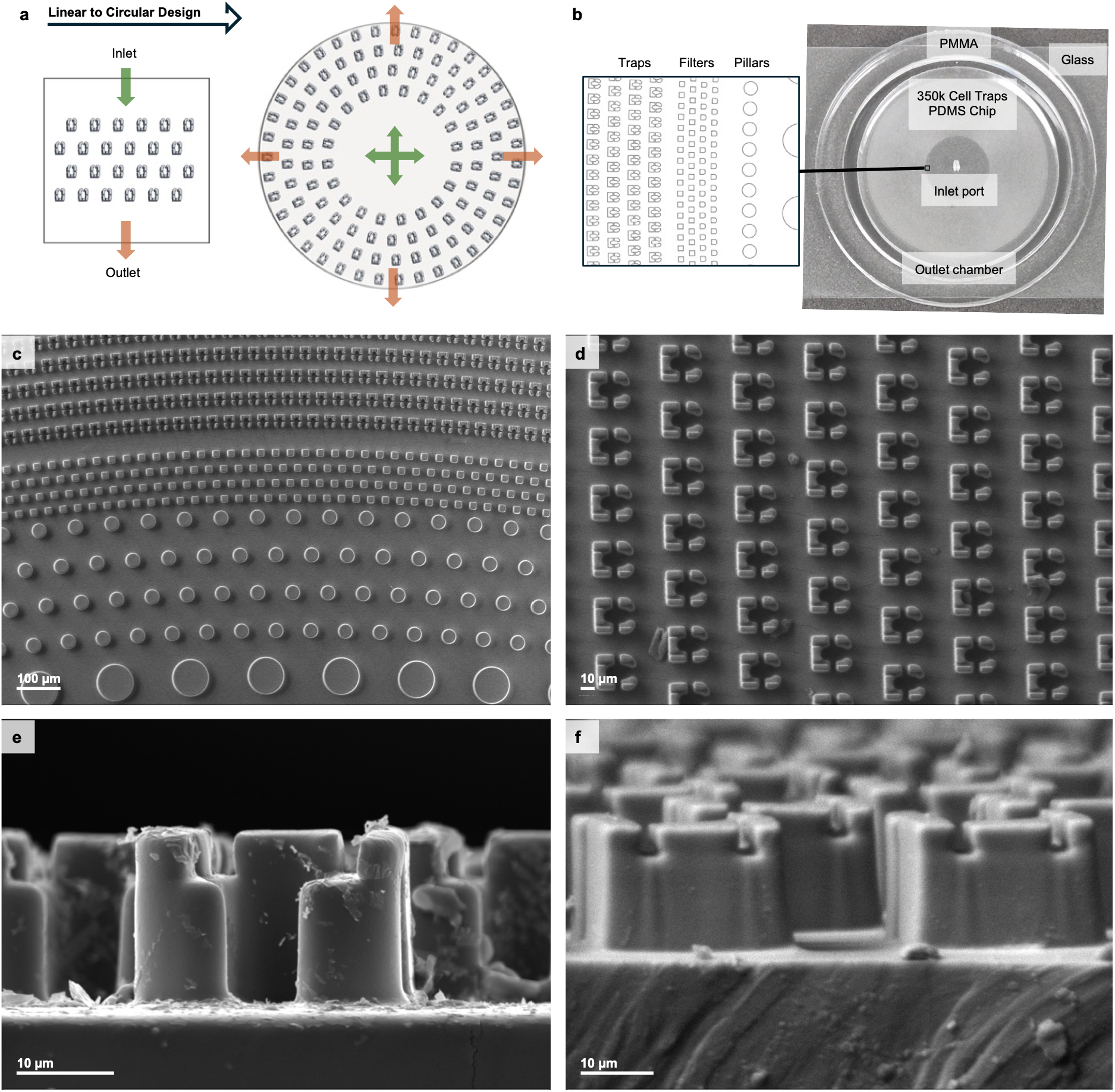
High-throughput cell pairing microfluidic chip design. a) Schematic representation of the transition from linear to circular trap design that allows the arrangement of larger number of traps. b) The image shows the final microfluidic device with its components: a PDMS chip (pillars, filters and 3D cell traps) with an inlet port followed by a waste chamber created around the chip with a PMMA ring assembled by PSA. c) SEM image showing the pillar arrays for filtering cells and the trap array for cell pairing and fusion. d) Top image showing the elements and dimensions of the 3D cell traps. f and e) Front and back image of the 3D cell trap showing its height.

The PDMS microfluidic chip with its components is shown in **Figure 1b**. The device has an inlet (central port) and liquid leaves through the external diameter of the chip. A chamber to contain and remove waste is formed by the space created between the chip and a PMMA ring attached to the glass slide by double-side adhesive. A set of pillars is placed immediately around the inlet to filter cells and to prevent collapse of the PDMS ceiling (**Figure 1c**). These pillars decrease in size towards the trap section, filtering cells progressively, and avoiding clogging. Downstream lies the cell-trapping area, formed by concentric rings around the inlet containing ~350,000 traps in total. Finally, a second array of pillars downstream of the trapping area provides structural stability to the chip and allows liquid to leave the device.

The cell-trapping area is densely packed with weir-based hydrodynamic traps arranged in a flow-through channel (**Figure 1d**), enabling deformability-based pairing of two sequentially loaded cell populations. The trap geometry (constriction and support pillars) and the deformability-based pairing protocol follow the microfluidic chip reported by Dura et al. ^[18]^ Each trap unit consists of a single-cell cup that arrests a single cell (the single-cell trap) connected through a narrow constriction to a downstream double-cell trap, together with support pillars at the sides and rear that sustain continuous perfusion once the trap is occupied. Because both cell types used in this study (immortalized mouse dendritic and tumor cells) have a mean diameter of 10 μm, the width of the single-cell trap was set to 10 μm and the length of the double-cell trap to 20 μm, so that exactly two cells are accommodated while minimizing the probability of retaining three or more. The constriction was set to 5 μm after empirical testing. Wider constrictions allowed multiple cells to enter the double-cell trap during the single-cell loading step, whereas narrower constrictions reduced single-cell capture efficiency. Lateral openings of 3-4 μm in the wall of the double-cell trap, together with the support pillars, route liquid through the trap rather than merely around it. This flow-through geometry retains captured cells against escape while permitting uninterrupted perfusion of the array after trapping, so that reagents can be exchanged across a fully occupied chip. This is a prerequisite for the timed PEG delivery described below. The support pillars also determine the fraction of local flow routed through the constriction and therefore the hydrodynamic force available to deform a captured cell into the double-cell trap. The whole 3D trap structure has a height of 17 μm (**Figure 1e** and **1f**), which both limits vertical overlap of cells and, together with the constriction, confines each cell so that transfer proceeds by controlled deformation rather than by bypass.

### 2.2 Automated chip priming and reagent exchange

The automated fluidic system uses a pressure-driven architecture with a flow switch to deliver reagents and cells to the chip (**Figure 2a**). It was assembled from Elveflow components and software together with laboratory-customized reservoirs, whose fabrication is described in Supplementary Information S4. A single pressure source drives liquid into the chip through a manifold that splits the applied air pressure among the reservoirs. Each reservoir is connected to a distribution valve (liquid manifold) that selects the liquid to be injected, which then passes through a thermal flow sensor used to monitor and regulate the volumetric flow rate. The system accommodates up to 11 reservoirs, and the injection sequence is executed from a computer through the Elveflow Smart Interface. The pressure, flow rate and duration applied at each step of chip priming, cell trapping, cell pairing, cell fusion and incubation are summarized in **Figure 2b**. The reported values are from the moment the valve is turned on. It is noteworthy that tube length from the liquid distribution valve to the chip inlet determines the delay between valve actuation and arrival of each solution at the trap array. For instance, with the 50 cm length used here this delay is up to 3 min (see Supplementary Information S5 and Figure S2), which depends on the properties of the sample. Because one pressure source supplies the driving force and the reservoirs are addressed through fluidic selectors, exchanging a reagent requires only replacing a tube, so a reagent change is a single tube swap rather than unloading, refilling and repriming a syringe. The flow sensor and selector are reusable elements in the flow path. Reusable microfluidic flow sensors of this type have been operated in an extracorporeal circuit in a rat model, ^[33]^ which demonstrates that sensors of this kind can be operated in an in vivo circuit. Nevertheless, sterilization and single-use qualification would need to be reconsidered for clinical manufacturing. The transparent chip and fluidic system are coupled to a fluorescence microscope to monitor cell pairing and fusion for subsequent image analysis.

**Figure 2.**
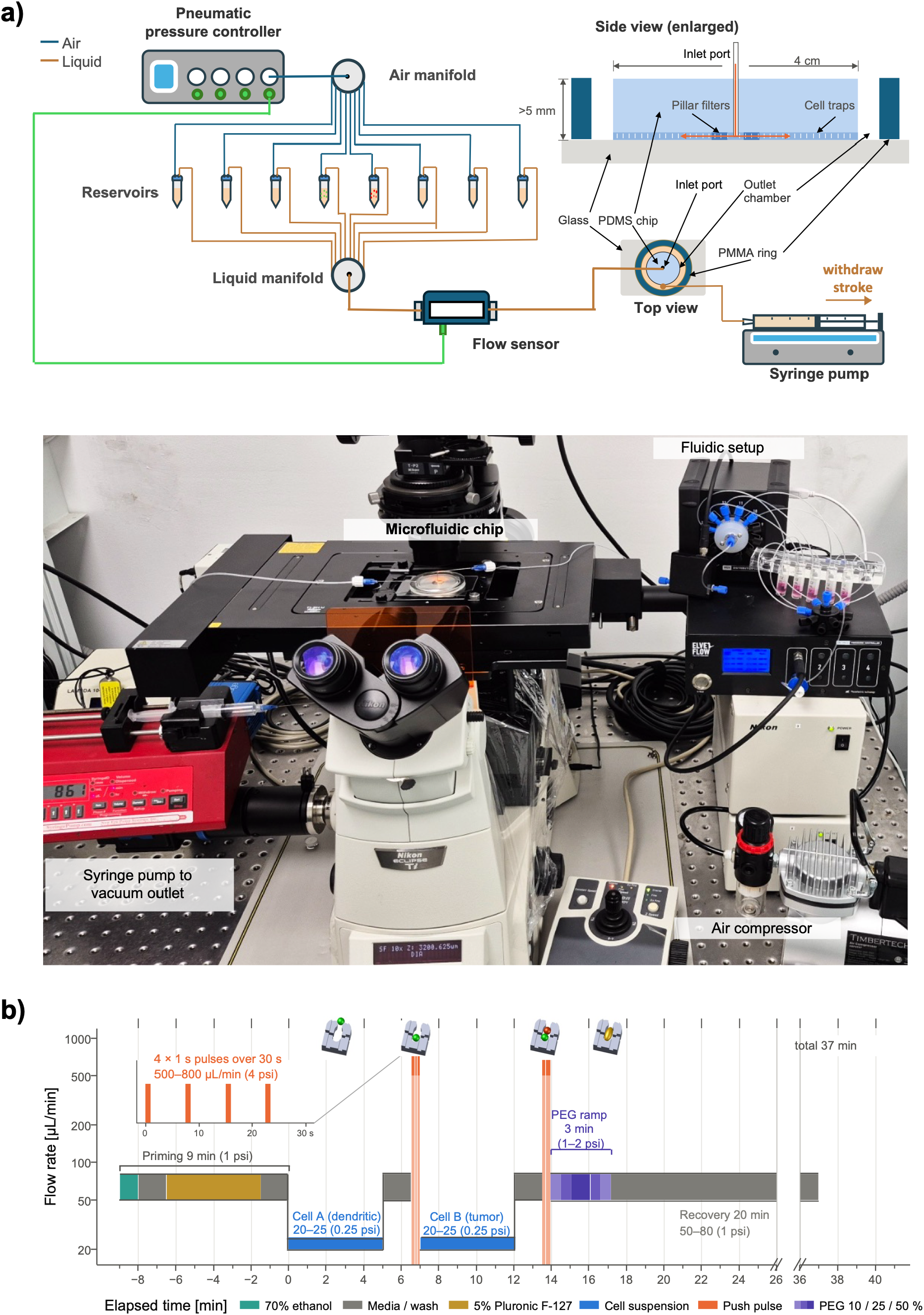
High-throughput cell pairing microfluidic platform for chemical cell fusion. a) Schematic representation and real photo of the entire platform for automated microfluidic cell pairing and fusion: the process is monitored by an inverted fluorescence microscope, the reagents are injected using a pneumatic pressure-based fluidic system driven by an air compressor, and a syringe pump removes waste. The chip side view is enlarged for better appreciation of the chip components. b) Automated chip priming, cell pairing process and optimized stepwise PEG-based protocol for cell fusion.

### 2.3 High-throughput cell pairing

We achieved cell pairing using the capture and trapping protocol shown in the upper part of **Figure 3A** and **Figure S1**. After cleaning and priming the tubing from the fluidic system as well as priming the chip with 75% ethanol, 5% Pluronic F-127, and cell media (see **Figure 1b**), the first cell population (dendritic cells) is injected at a concentration of (8×10^6^ cells/mL) and low flow rate (20-25 μL/min). The low flow rate ensures capture at the front opening of the single-cell traps and the selected cell concentration allows the feeding of the entire chip while avoids clogging. After 5 min, once the single-cell traps are saturated, the switch turns to media, and the flow rate is raised for 90 s to remove uncaptured cells. At this stage the flow rate must not exceed 100 μL/min, above which cells risk being deformed and transferred into the double-cell traps permanently, particularly in the rings closest to the center. Four pulses of high flow (500-800 μL/min) were then applied to deform and transfer the captured cells from the single-cell traps into the double-cell traps (**Figure 3a**). The sequence is repeated with the second population (tumor cells) to complete pairing. Large-area images were then acquired to calculate trapping and pairing efficiency by a MATLAB code using the Image Processing Toolbox (**Figure 3a, bottom**). This optimized protocol achieved a pairing efficiency of approximately 75% (**Figure 3b**), defined as the fraction of traps containing at least one tumor cell and one dendritic cell and no more than three cells in total. Approximately 40% or ~140k traps contained exactly one tumor cell and one dendritic cell. In addition, **Figure S3** in Supplementary Information S6 shows that the pairing efficiency measured across the radius from fluorescence tiled images is ~73% and ~64% in the innermost and outermost rings, respectively. This proved that the operation flow rates used here do not influence significantly the pairing efficiency across the radius.

**Figure 3.**
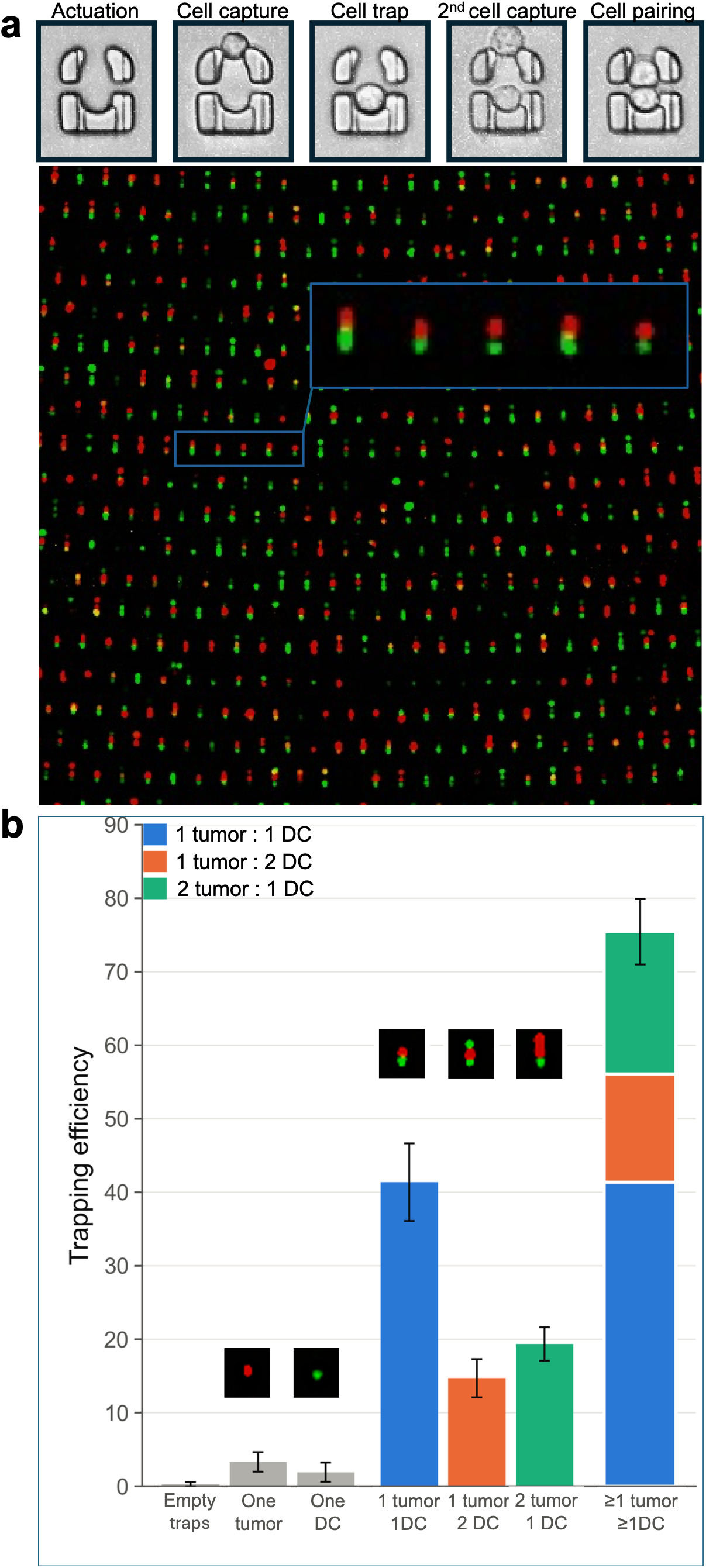
High-throughput cell pairing results using tumor cells and dendritic cells (DC). a) Cell trapping and pairing process (top) and small portion of a multichannel fluorescent large image showing the cell pairs (bottom). b) Trapping efficiency calculated by a Matlab code. Each value is the mean of 3 independent experiments. Tumor and dendritic cells are immortalized mouse C1498-RFP and MutuDC1940-GFP cell lines, respectively.

### 2.4. Stepwise PEG delivery optimizes chemical fusion

After the automation of high-throughput cell pairing, we optimized the sensitive and toxic process of PEG-mediated chemical fusion of mouse dendritic and tumor cells. PEG drives membrane apposition and fusion but is cytotoxic through several concurrent mechanisms. At fusogenic concentrations, it withdraws water from the intermembrane space and imposes osmotic and curvature stress on the bilayer. This crowds the extracellular macromolecular environment, dehydrating and destabilizing the membrane surface, and therefore sustained exposure perturbs bilayer integrity and lowers post-fusion viability.^[30,34]^ Because these effects scale with both concentration and exposure time, we reasoned that delivering PEG as a step sequence rather than a single step change would reach fusogenic conditions while limiting the abruptness of the osmotic transition. We therefore compared three on chip stepwise delivery protocols: a step-up sequence (media, 10%, 25%, 50% PEG, media); a step-down sequence (media, 50%, 25%, 10% PEG,media); and a step-up-and-down sequence (media, 10%, 25%, 50%, 25%, 10% PEG, media). A single 50% PEG 1500 exposure was run on chip alongside the step-based protocols as control. The conventional benchtop protocol using 50% PEG 1500 in an Eppendorf tube is reported separately (**Figure 4c and d**) as an additional control. The step-up and the step-down sequences are therefore not a control in the sense of an untreated reference, but partial-protocol comparators that isolate the contribution of the ascending and descending arm of the step-up-and-down PEG profile. It is important to note that the PEG profiles were selected experimentally, and it was found that using 1 min of critical 50% PEG exposure on chip was enough to reach adequate cell fusion while keeping cell viability ≥ 90%. In each case, the PEG sequence was followed by injection of fresh media at 50-80 μL/min for 20 min, which simultaneously washes residual PEG out of the occupied array and allows membrane rearrangement and cell recovery before collection.

**Figure 4.**
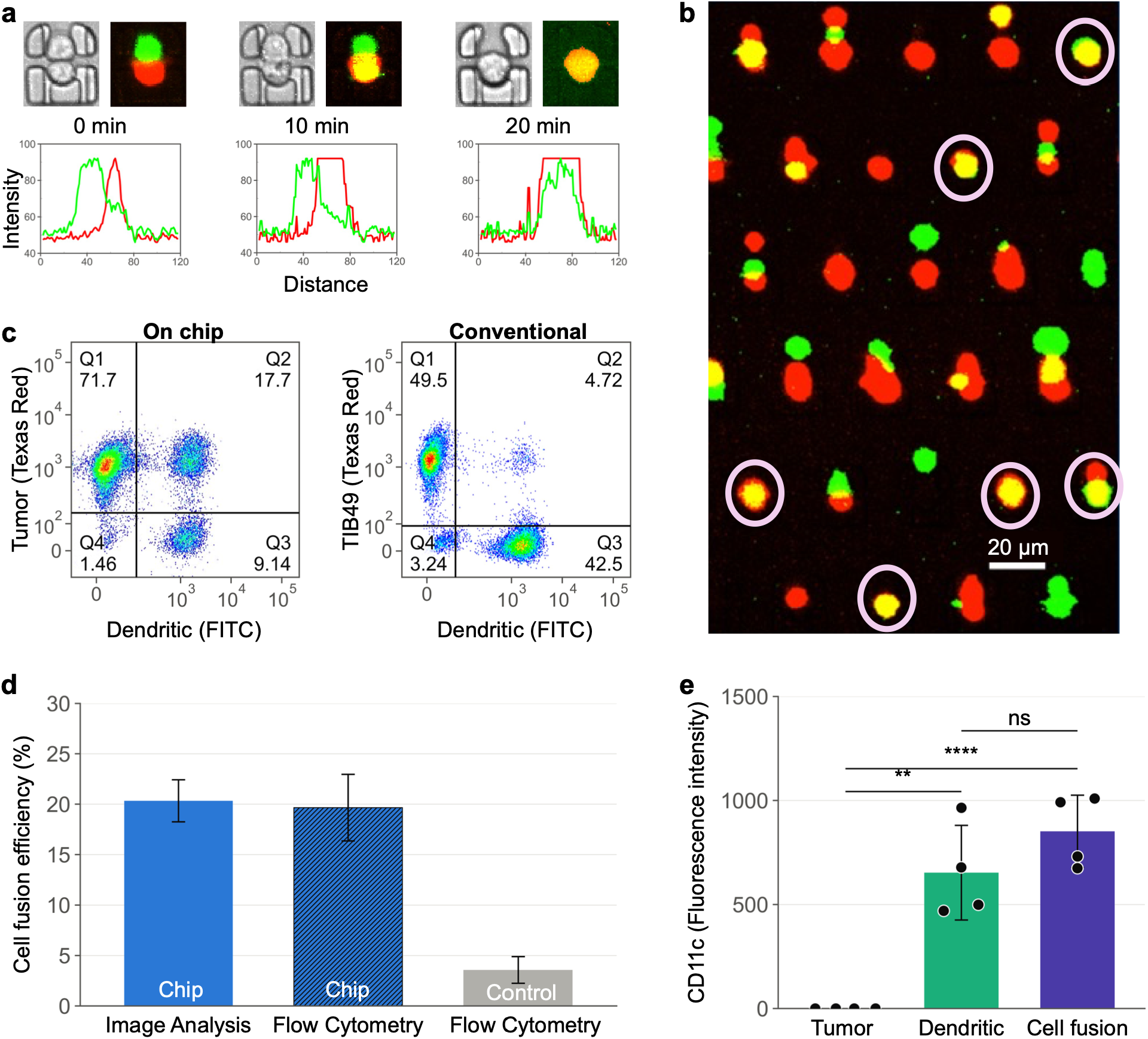
Demonstration of PEG-based cell fusion of tumor (C1498-RFC) and dendritic (MutuDC1940-GFP) cells in the automated microfluidic platform. a) Zoomed view of a representative region from tile images showing microfluidic traps exhibiting colocalization (yellowish color) of both channels (pink circles), resulting from the merge of red and green fluorescence emitted by cells undergoing fusion. b) Time-lapse monitoring of the fusion process where cell membrane rearrangement and merging of fluorescence signals are observed through peak intensity plots. c) Flow cytometry-based fluorescence analysis of fusion cells generated in the automated microfluidic platform, combining dendritic cells and tumor cells signals detected in the FITC and Texas Red channels, respectively. The Q2 quadrant identifies the double-positive (GFP^+^/RFP^+^) population, corresponding to successfully fused cells in the automated microfluidic platform and conventional method. d) Comparison of the cell fusion efficiencies using both image analysis and flow cytometry analysis. e) Analysis of CD11c marker expression in dendritic cells, tumor cells, and fused cells.

The step-up-and-down protocol preserves viability above 90% and yields a fusion efficiency of ~18% by image analysis in the primary run. This result agrees with the flow-cytometry mean reported below and is also consistent with the hypothesis that a gradual rise in PEG concentration avoids a sudden osmolarity change and thereby limits membrane damage. Efficiencies for step-up and step-down PEG sequences from image analysis were 8% and 6%, respectively, which shows that the complete step-up-and-down profile improves fusion by at least two-fold relative to the other partial comparators.

All the steps described above (cell loading, pairing, PEG delivery and washout) were performed at room temperature on the microscope stage with no active temperature control applied. Therefore, the efficiencies reported may underestimate those achievable with active temperature control, and adding a stage-top incubator is a direct route to improvement. **Figure 4a** shows a region extracted from a large-area fluorescence image acquired with a 10x objective at an exposure of 900 ms per channel, in which hybrid cells appear as single cells with colocalization of both channels (yellow). Images of this kind were used to calculate cell fusion efficiency by a custom script in MATLAB where cell fusion is set as the colocalization of the maximum peak intensity inside a trap from both fluorescence channels. To corroborate the fusion efficiency obtained by image analysis and to characterize the resulting hybrids, we analyzed all cells recovered from the chip by flow cytometry after the fusion process and 24 h of incubation. The incubation period allows the hybrids to recover from the stress imposed by pairing and PEG exposure and ensures adequate viability (>90%). **Figure 4c** shows that hybrid cells appear as a dual-fluorescent population (quadrant Q2) arising from the merging of cytoplasmic content from both partners. Flow cytometry gave a fusion efficiency of 17.7% for the step-up-and-down PEG profile, in close agreement with the image-analysis value. Two further independent replicates (Supplementary Information S7, **Figure S4**) gave 18.4% and 23.4%, giving a mean of 20% across three independent experiments, i.e., range 17.7-23.4%. These results corroborate the corresponding image-analysis results. The obtained efficiency is more than three-fold higher than that obtained with the conventional protocol performed in an Eppendorf tube (4.72%; Figure 4c and 4d). **Figure S4b-d** additionally shows flow cytometry for the step-up, the step-down, and the 50% PEG single-step protocols run on chip, which yielded 8.17%, 7.10% and 6.17%, indicating that the complete step-up-and-down profile improves fusion by at least two-fold relative to any of the partial or single-step comparators.

Until prior to chemical cell fusion, paired and single cells can be collected without disassembling the device, by injecting media through the outer rim outlet while applying vacuum at the central inlet to reverse the flow direction through the array (<u>Supplementary Video1</u>). For data presented, cells were recovered by detaching the PDMS layer from the glass slide and rinsing the exposed trap array with media. Because recovery requires disassembly of the device, the correspondence between an individual trap and a recovered cell is not preserved, and all downstream analyses are therefore population-level. The recovery efficiency, defined as the proportion of cells present in the array that were retrieved for downstream analysis, was 80%. This value sets the practical yield of the platform of the approximately 55,000-60,000 hybrids formed on chip, approximately 50,000 are available for off-chip characterization.

We additionally assessed retention of the dendritic-cell surface marker CD11c in the fused population (**Figure 4e**). CD11c is expressed by dendritic cells but not by tumor cells, so its presence in a dual-fluorescent event is consistent with incorporation of dendritic-cell membrane into the hybrid. **Figure 4e** shows that tumor cells are negative for CD11c, whereas dendritic cells and fused cells are positive. This is a single-marker surface-expression analysis rather than phenotyping. One lineage marker establishes that dendritic-cell membrane is retained after fusion, but it does not establish the maturation or functional state of the hybrid, which would require a panel including co-stimulatory and antigen-presentation markers (for example CD80, CD86 and MHC class II) together with a functional readout. Retention of CD11c is nevertheless a necessary precondition for the antigen-presenting function on which the dendritic-tumor hybrid vaccine concept depends [30], [31]. Taken together, these results show that the automated microfluidic platform generates approximately 50,000 viable fused cells per chip in a dendritictumor cell model.

## 3. Discussion

The weir-based cell traps enable dynamic monitoring and manipulation of single cells and cell-cell interactions, including cell-cell fusion for therapies. In this work, the radial arrangement of cell traps followed by automation (**Figure 2**) are presented as a methodology for high-throughput cell trapping and pairing operations. The automation is necessary for reproducible serial fluidic steps (cleaning and priming of the tubing, chip priming, sample preparation and final cell fusion) all of which required up to 159 minutes (**Figure 2b** and **S2**). We demonstrated that the radial decline of samples and reagents flow rate along the increasing radius is within operational range of the presented cell-lines and applications. As mentioned in a patent application,^[32]^ minimizing the flow-rate difference between the innermost and outermost trap rings can be further addressed by reducing the number of the cell trap rings and instead increasing the radius of the cell trap zone. Alternatively, the constriction sizes between single cell and double cell trap can be graded as a function of radius to make the best use of the available space in the chip. The developed automated high-throughput platform reproducibly achieved ~75% cell pairing or trap occupancy by at least one cell of each type i.e., 250,000 of total 350,000 traps (**Figure 3b**). Approximately 140,000 of those are one-to-one pairs of one tumor and one immature dendritic cell. The one-to-one fraction (~40%) is lower than previously hydrodynamic microfluidic devices of similar mechanism ^[3,18,24]^ but it is obtained at a significantly larger throughput (at 10^5^ scale). This is of particular importance for robust process optimization, adequate cell production and downstream multiplexed analysis e.g., dynamic image microscopy on the chip and flow cytometry off them chip. The platform was subsequently used to optimize the relatively short but highly sensitive/toxic process of chemical cell fusion of immortalized tumor (C1498-RFP) and dendritic (MutuDC1940-GFP) cell lines. These cells were used as a model system because of their potential for personalized cancer immunotherapy. Stepwise PEG protocols were implemented based on the hypothesis that, beyond exposure time, controlling osmotic stress from toxic PEG can affect the fusion efficiency and cell viability. Minimal flow rates were selected experimentally to ensure delivery and removal of PEG across all traps, and therefore the effect of flow rate on fusion is not a studied variable. In addition, as part of automation for the development of a micrototal analysis system, fusion events in the chip were quantified by image analysis via a custom MATLAB scrip and validated by flow cytometry. The strategy to minimize the risk of osmotic shock was to gradually change PEG concentration with two additional steps of 10% and 25% before or/and after the critical high dosage of 50%. These stepwise protocols were compared to two control tests, the single 50% PEG step on chip and the conventional method performed in tubes. The on-chip control test shown in Figure S4d results in a higher fusion efficiency and adequate viability when using only 1 min of exposure to the high concentrated reagent (e.g., 50% PEG), which is at least 33% shorter than commonly exposure duration used in conventional PEG-based protocols. While fusion improves at both rap-up and step-down (**Figure S4b** and **S4c**), under the optimized step-up-and-down protocol the cell fusion process yielded an efficiency of ~20% vs. ~6% at single the 50% PEG profiles on chip and ~5% for the conventional control (**Figure 4b** and **S4d**). Therefore, the platform generated approximately 50,000 viable collected fused cells per chip that is adequate for downstream off the chip analysis and use. In addition, the characterization of CD11c marker on the hybrid cell population as shown in **Figure 4e** indicates the correct integration of dendritic cells in the cell fusions. Overall, automated high-throughput methodology, beyond facilitating personalized cancer vaccine generation research presented here, can be adapted to several existing microfluidic cell engineering to enable from dynamic in-vitro processes to clinical sample preparation and testing. Finally, the design methodology is aligned towards integration into the centrifugal microfluidic system to take advantage of the pseudo forces from the chip spinning.

## 4. Conclusion

This work establishes an automated high-throughput microfluidic platform for cell pairing and chemical fusion. A radially symmetric chip arranges ~350,000 deformability-based cell traps in concentric circular arrays and receives all reagents through a single central port. The platform co-occupies approximately 75% of the traps (~250,000) with at least one cell of each type, and approximately 140,000 of these traps hold exactly one tumor cell and one dendritic cell. The resulting cell pairing throughput per chip exceeds that of previously reported hydrodynamic pairing arrays by more than one order of magnitude. A symmetric step-up-and-down PEG profile results in fusion efficiency of approximately 20% (range 17.7 to 23.4%, n = 3) with viability above 90%. Therefore, each chip generates at least 50,000 hybrid cells. Fusion events were quantified by image analysis and validated by flow cytometry. Retention of CD11c biomarker indicates that hybrid cells preserve the dendritic-cell phenotype after cell fusion. Automation governs both the PEG concentration profile and the timing of reagent exchange, and the protocol therefore remains reproducible across runs. The underlying design strategy used here is not specific to cell fusion and is possible to transfer to other cell applications. The cell fusion throughput of this platform provides sufficient hybrid cells for downstream immunological evaluation and further cancer-vaccine development.

## 5. Experimental Section

### Device fabrication

Master molds were produced by two-layer photolithography on silicon wafers using negative photoresists SU-8 2005 and SU-8 2010 (MicroChem), giving a total feature height of 17 μm. Devices were cast in PDMS (Sylgard 184) at a base-to-curing-agent ratio of 10/1 and cured at 80 °C for 2 h. Cured slabs were peeled from the mold, cut to size and punched at the inlet and outlet, then bonded to glass slides by oxygen plasma treatment at 100 W for 60 s, followed by heating at 80 °C for 5 min to consolidate the bond. Also, to avoid breaking the master mold, copies were fabricated using polyurethane resin (Smoth-Cast™ Onix™ Slow).

### Cell culture

The MutuDC1940-GFP cell line (immortalized mouse dendritic cells) was obtained from Applied Biological Materials Inc. (Vancouver, Canada). The cells were cultured under the supplier’s recommended conditions: Prigrow V medium supplemented with 4 mM Glutamax, 10% fetal bovine serum (FBS), 1% sodium bicarbonate solution, 50 µM β-mercaptoethanol, 10 mM HEPES, and 1% penicillin/streptomycin. These cells stably express GFP genes and are CD11c^high^. The C1498 murine leukemia cell line (ATCC TIB49, immortalized tumor cells) was obtained from the American Type Culture Collection (ATCC). The cells were cultured in RPMI 1640 medium supplemented with 10% fetal bovine serum (FBS). Both cell lines were maintained at 37 °C in a humidified CO2 incubator under a 5% CO2 atmosphere. Before performing cell pairing and fusion process, the viability of all the cell samples were above 90% measured by standard trypan blues testing.

### Generation of Stably Expressing TIB49-RFP Cell Lines

We developed a fluorescent C1498 cell lines (RFP and GFP) by electroporation to enable robust analysis of cell pairs and fusions using fluorescence microscopy and flow cytometry. To generate these fluorescent cell lines, we designed two plasmids carrying GFP and RFP gene under the control of the EF-1α promoter. These plasmids also include the puromycin resistance gene to ensure stable expression in the cells. Detailed plasmid maps are provided in the supplementary material (Supplementary Information S8). Transfection was performed using the Gene Pulser Xcell system (Bio-Rad, California, USA) and its corresponding electroporation cuvettes. We resuspended 250,000 TIB49 wild cells in phosphate-buffered saline (PBS), and 5 μg of plasmid DNA was added. The process was carried out using a single pulse at 200V for 10 ms. Finally, transfected cultures were then selected with puromycin for three weeks, and single-cell sorting was performed using a Cytoflex SRT flow cytometer (Beckman Coulter, California, USA).

### Cell trapping pairing

Prior to using the pressure-based fluidic system, it was subjected to a cleaning process using 70% ethanol, PBS and 10% bleach followed by tube priming with 70% ethanol, 5% Pluronic F-127, and cell media (see Supplementary Information S5 for details). Cell trapping is performed after priming the device with 70% ethanol, which displaces air from the channels and trap array and thereby removes bubbles, followed by injection of 0.2% Pluronic F-127 to prevent cell adhesion to the microfluidic walls. The trapping and pairing sequence consists of loading 100 μL of the first cell suspension at 8×10^6^ cells/mL for 4 min at 0.25 Psi (20-25 μL/min) to capture cells at the openings of the single-cell traps. Fresh complete media is then injected to remove excess cells (1 min at 1 Psi, 50-80 μL/min). Four pulses at 4 Psi (500-800 μL/min) are subsequently applied over 1 min to deform the captured cells through the constriction into the double-cell traps. The sequence is repeated with the second cell suspension to complete pairing. All steps are performed at room temperature. Brightfield and fluorescence images of four large regions (10-20 fields each, 10x objective) are then acquired on an automated inverted microscope (Nikon Eclipse Ti). The images were analyzed in MATLAB 2026b using a custom scrip and the image processing toolbox to calculate pairing efficiency. The code quantify efficiency by segmenting individual microwell traps from brightfield images, counting dendritic and tumor cells within each trap via fluorescence peak detection to finally calculate the percentage of traps with different tumor-dendritic occupancy combinations across the segmented trap area in the tile images.

### Cell Fusion

After the cell pairing procedure and the associated image acquisition, when each trap contains at least one cell of each type, PEG-based fusion is performed in the device. Preliminary tests were carried out to optimize the protocol with respect to fusion efficiency and viability. The optimized process consists of stepwise increases in the concentration of the injected PEG solutions followed by symmetric decreases, which reduces cell damage caused by abrupt changes in osmolarity. The order of the injected solutions is: 10% PEG1500 (30 Psi, 100 s), 25% PEG1500 (30 Psi, 90 s), 50% PEG1500 (2 Psi, 60 s), 25% PEG1500 (1 Psi, 30 s), 10% PEG1500 (1 Psi, 30 s). Cell media is then injected at 1 Psi for 20 min to allow recovery and membrane rearrangement. All steps are performed at room temperature on the microscope stage. During fusion, a region is monitored to follow progress, and images are acquired before and after fusion to quantify efficiency in MATLAB. Also, cell fusion efficiency is calculated by measuring the spatial coincidence between the dendritic and tumor fluorescence intensity peaks within each trap, with peak overlap ≥ 90% of the trap’s spatial footprint scored as fusion positive. After fusion, the PDMS chip is carefully detached from the glass slide; detachment without shearing the array is possible because the design contains no continuous bonded wall enclosing the trap region, so the array is exposed uniformly as the layers separate. Cells are then collected by rinsing with media and incubated at 37 °C for 24 h. The recovery efficiency under these conditions is 80%. To limit contamination risk during detachment and rinsing, these steps are carried out inside a biosafety cabinet.

### Flow cytometry

Twenty-four hours after the fusion procedure, the cells were resuspended in PBS and efficiency of cell fusion was calculated using a celesta flow cytometer equipped with 405 nm, 488 nm, and 640 nm lasers (BD Biosciences, New Jersey, USA). All the cell samples had at least 90% of viability after incubation. Additionally, dendritic surface markers in cell fusions were analyzed by flow cytometric immunostaining. A primary anti-CD11c antibody (Cat. 14-0114-82, Thermo Fisher Scientific, Massachusetts, U.S.A.) and a secondary Goat Anti-Armenian Hamster IgG Alexa Fluor® 405 antibody (Cat. ab175680, Abcam Limited, Cambridge, United Kingdom) were used for staining. Briefly, 150,000 cells were incubated with the primary anti-CD11c antibody for 30 minutes on ice. Following incubation, the cells were washed three times with PBS and then stained with the secondary antibody for an additional 30 minutes on ice. Afterward, the cells were washed three more times with PBS before analysis.

## Acknowledgements

The authors acknowledge the financial support provided by NIH (R21 CA259840), the Tecnologico de Monterrey-MIT program, and the Federico Baur Endowed Research Chair from Tecnologico de Monterrey. The authors thank the FEMSA Biotech Center for their support in providing workspace. The authors also thank the scholarship provided by SECIHTI and Tecnologico de Monterrey.

## Conflict of Interest

The authors declare no conflict of interest.

## Data Availability Statement

The data that support the findings of this study are available in the article and the supplementary materials of this article.

## Notes

### Competing Interest Statement

The authors have declared no competing interest.

