## Supplementary Information for "Automated High-Throughput Microfluidic Platform for Cell Pairing and Chemical Fusion"

**S1. Array-construction methodology**

The array is built by a spoke-and-circular-array workflow that turns a single drawn unit into the full device and, in the same step, enforces the deterministic cascade that drives capture. First, a single radial spoke of six to eight single-cell traps is drawn, with alternate traps shifted circumferentially so that, once arrayed, each ring's openings register behind the inter-trap gaps of the ring immediately inward of it. For example, in a spoke of six to eight traps, every second trap can be rotated by 360° / (twice N(r)), or 180° × (pθ / (2π r)), prior to applying simple radial replications in the second step. For radial replication, the spoke is arrayed across the radial span $r_1$ to $r_2$ to seed every concentric ring position. This is also the appropriate stage to introduce radius-graded trap constrictions if desired as a flow-compensation option, before applying circular array command of CAD software e.g., AutoCAD or SolidWorks. In the next step, each spoke from radial replications is circular arrayed about the central inlet. Since the number of traps N(r) is re-incremented for the next group as the circumference grows, recounting them is necessary.

Because each circular-array command fixes a single integer N(r) for one group of rings, the realized circumferential gap is not exactly constant but drifts with radius. The radial span of each fixed- N(r) group is therefore capped to keep the gap within the design band, which is the origin of the six-to-eight-trap spoke length. Openings are registered behind the upstream gaps by offsetting consecutive rings by half the angular pitch. Because N re-increments between groups, exact registration cannot be held across a group boundary. This is a commensurability limit intrinsic to building rings from identical units, which slightly lowers pairing yield but does not affect the pressure–flow behavior.

Two features make this a transferable high-throughput methodology rather than a one-off layout. First, the array commands (one linear array for radial groups, and circular arrays for each with numbering N(r) for each radius) expand a single drawn spoke into the entire array. Second, the half-pitch stagger lets a single radial pass self-fill successive ring, raising fill fraction without active control. The recipe is geometry-agnostic, meaning that, substituting a different unit trap ports any planar hydrodynamic-trap design onto concentric rings.

### S2. Fabrication and scale-up notes

Master fabrication uses a two-layer SU-8 process (support pillars in the first layer and trap features in the second) patterned by direct-write laser lithography, after which the master is silanized and used to mold PDMS chips that are oxygen-plasma bonded to glass (full recipe in Methods). Two practical points govern how large the disc can be made i.e., Layer alignment and tolerance to trap-size error.

The two lithography layers must be aligned before the second exposure. Over a large disc, a small angular misalignment that is barely visible near the center is amplified toward the rim, where it can render rim traps non-functional and, more importantly, distort the radial symmetry that gives the array its uniform flow in every direction. We therefore place several registration marks of graded size (down to ≈5 µm, which defines the finest achievable alignment) at a minimum of three locations around the outer rim rather than near the center, where sub-visual errors would scale outward undetected. Good rim alignment preserves both trap function and the homogeneous radial flow on which cell pairing depends and is central both to scale-up and to reproducible fabrication.

The device is forgiving one class of fabrication error and sensitive to the other. If the trap openings come out larger than designed, capture and pairing still proceed, only at a lower flow rate. If the constriction comes out smaller than designed, the higher flow needed to seat cells and to transfer them from single to double traps can become excessive on the inner rings, where the local velocity is highest, so that cells may escape, be squeezed out of their traps, or be damaged. Erring toward a slightly larger constriction is therefore the safer manufacturing target.

### S3. Trials on careful manual one-to-one cell pairing process

Prior to automation, the feasibility of the system for cell pairing was tested with two syringe pumps and manual processing for disconnection of the inlet to load TIB-49 cells (tumor cells) of different fluorescence colors (RFP and GFP) into the injection needle. To prevent accidental bubble injection into the chip, it is crucial for the injection needle at the center to always discharge liquid when inserted or withdrawn from the chip. Therefore, as well it is essential to ensure the absence of air bubbles in the three syringes for the priming step, which contain filtered 70% alcohol, 2-5% Pluronic F-127 solution, and cell media.

The first priming step is the alcohol injection to wet the chip without the risk of air bubble traps between or inside cell traps is a rapid (five to fifteen seconds) but sensitive initial step as a single bubble that often later expands can ruin the homogeneous liquid and sample flow in the chip.

While the handheld alcohol syringe is discharging, first a drop is formed on the inlet surface while inserting the needle into the inlet. This step requires gentle insertion of the needle and liquid injection (touch). Often, especially within 1-2 hours of plasma binding, the solution's tendency to slowly drive through channels is visually evident, therefore it’s highly suggested to not vertically push syringe against hard against the glass surface. When alcohol reaches the edges of the chip either by gentile press or capillary flow, for adequate flow in order for the alcohol to exits the output ring press the syringe with slightly higher and gradually increasing force (patiently) and once the alcohol in the edges are visible, slowly and gently withdraw the needle while still the alcohol is discharging.

The next step is washing ethanol before Pluronic injection. Pluronic (e.g., F-127), a triblock copolymer (PEO-PPO-PEO), adsorbs onto hydrophobic surfaces like PDMS in microfluidic chips via hydrophobic PPO block anchoring, creating a hydrophilic PEO brush that prevents cell adhesion by repelling proteins and cells [35]. Some studies suggest brief 70% ethanol flushes for sterilization or preparation without noting issues [36], and coatings show stability in aqueous environments for weeks [37]. However, from our experience, injecting even low-concentration Pluronic F-127 (e.g., 0.1-2 wt.%) into a microfluidic chip after 70% ethanol without washing is suboptimal. The mixture of ethanol and Pluronic F127 creates a high risk of gel-like residuals that lead to uneven surface modification, channel blockages, and overall process failure. Scientifically, residual ethanol forms mixing gradients (e.g., 10-30 wt.% ethanol-water), which dehydrate the PPO blocks and trigger micellar aggregation into viscous clusters through mechanisms like PEO chain bridging or depletion flocculation [38]. While low concentrations remain untested in the literature, our experiments reveal similar residuals, likely due to shifted critical micelle concentration/temperature (CMC/CMT) and localized precipitation. Therefore, washing ethanol with PBS or cell media is essential before Pluronic injection and a 30-min incubation to ensure a stable, uniform coating. While the media syringe is dispensing liquid (over the alcohol droplet), gently insert it into the chip. The pinkish/reddish liquid should become obvious at the outer rim of the chip or the outlet. At this stage, gently wipe out the mixture of alcohol and media before removing the dispensing media syringe’s needle, so that a small droplet containing only media remains on the chip's center/surface.

After ethanol washing, and similar to the previous step, while Pluronic is slowly discharging, insert the needle and gently push the solution until the transparent liquid at the outlet of the chip obviously dilutes the red/pinkish color of the cell media. Then, gently remove the needle and let the Pluronic-loaded chip with the media-Pluronic droplet on the surface and incubate inside a closed Petri dish for at least half an hour.

Before using, wash the Pluronic with cell media and before removing the syringe wipe-out surface liquid and after letting a media droplet remain on the surface and liquid around the output of the chip to prevent the risk of evaporation and bubble formation from possible delays or errors, including microscopy light/lamp.

For the cell pairing step, the system works best with fresh cells, as fresh as possible. The thirty-minute Pluronic incubation is a good time frame to prepare the cells. For this manual procedure, having two syringe pumps is highly suggested. A high cell concentration in a small volume works best, although it comes with a high risk of cell adhesion if viability is below 95% but easier washing steps. A volume of 1 million cells in 100 microliters or 1 million cells per 60 microliters can fit into the injection needle connected to the syringe pump that is connected to cell media. This is of advantage as cell residuals often are formed in junctions in external fluidic networks. Therefore, the fresh cells are filtered with a 10-micron strainer at high concentration. For instance, a high concentration is formed as 150 microliters of up to 10,000,000 cells per ml solution or 100 microliters of 16,700,000 TIB-49 RFP-cells/ml solution. The injection needle of syringe pump is then inserted into the solution, and gently (30-50 microliters per minute) the solution is uptake into the needle. While still the needle is dipped into the solution, the pump is switched to 10-20 microliters per minute push mode. This prevents the uptake of air when removing the needle from the solution. Wait for seconds until a small droplet is formed on the needle tip to assure cell discharge before gently inserting it into the chip that contains a droplet of media on the center to prevent evaporation and bubble formation in center. Once the cells have reached the cell trap zone, the flow rate is elevated to 60 to a maximum of 80 microliters per minute, and the cell-capturing step occurs in less than a minute (**Figure S1**). Afterwards, the flow rate is increased for washing excess cells for up to 2 minutes. The flow rate then is reduced to 20 to 40 microliters for the cell trapping step with the second pump to prevent liquid backflow due to high liquid flow. The second pump is set to 2 to 4 ml/min per followed by applying several pulses (e.g., 2 or 6). The shear force from the high flow rate deforms the cells and pushes them into cell traps. Absence of the flow (e.g., 10- 20 microliter/min) can result in liquid back flow and cells entering the chip from outlet towards disc center which may cause both contamination and extra wash step. While leaving the flow at as low as 10-20 µL/min, second cells (TIB-49 GFP cells) were filtered with 10-µm strainer, and the entire process was repeated to accomplish cell pairing.

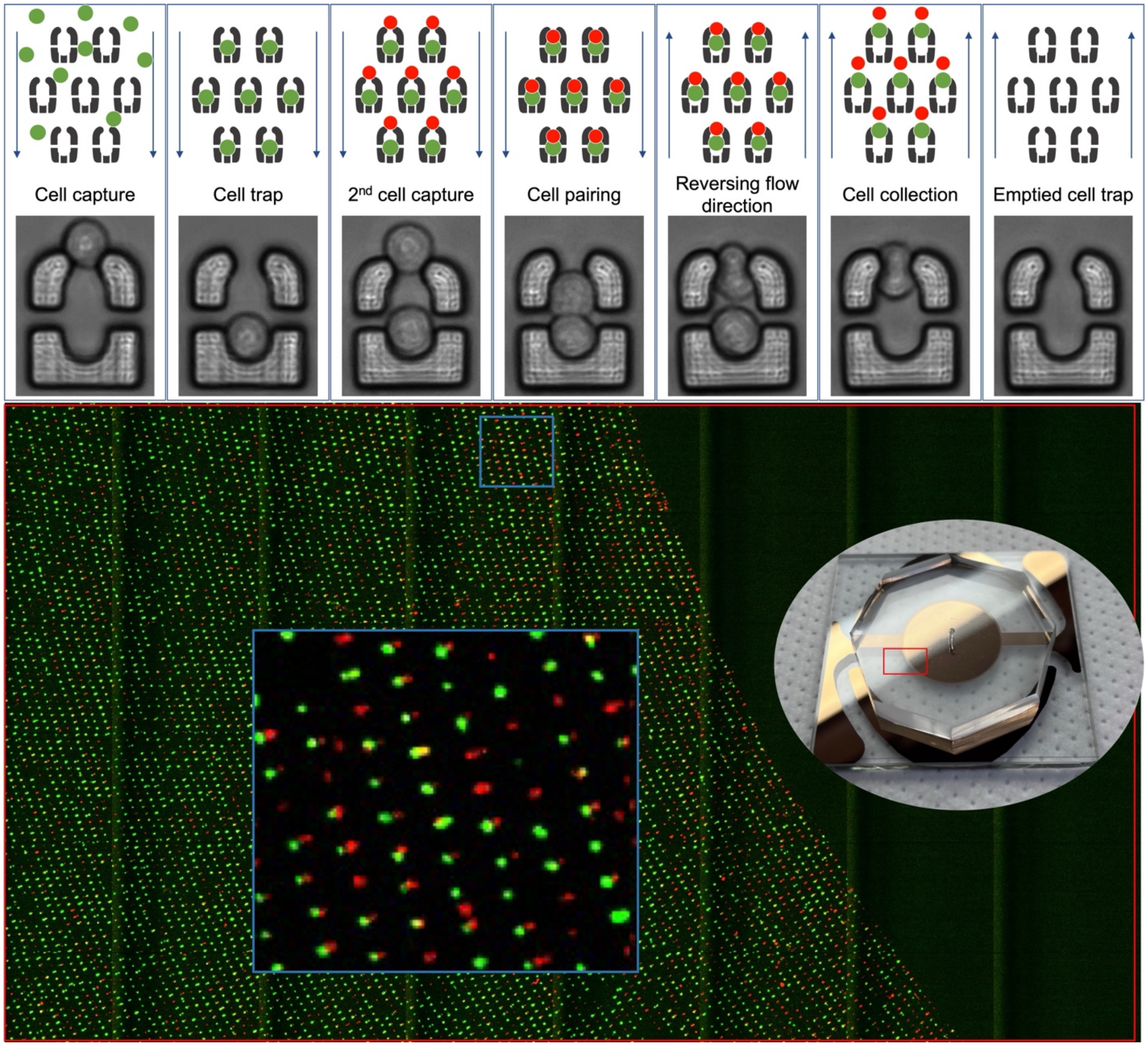


**Figure S1**. Manual one-to-one cell pairing process as a proof-of-concept of the radial chip functionality. The top sequence is the trapping C1498 GFP and RFP cells in the microfluidic chip, followed by their collection via revering the direction of flow. The bottom part is an example of high-throughput microscopy imaging from an area of the chip.

**S4. Additional details of the automated microfluidic platform for cell pairing and fusion**

Reservoirs were built from conventional microcentrifuge and conical tubes fitted with cast PDMS caps perforated to accept inlet and outlet tubing. The caps were molded singly or as strips in a flat mold and punched with a biopsy punch to allow for standard tubes to be adapted for pneumatic pumping at negligible cost.

**S5. Tubing cleaning and priming, and preliminary actuation.**

The pressure-driven fluidic system (tubing, reservoirs, flow sensor, and liquid distribution valve) was subjected to cleaning and priming process before each experiment remove particles, avoid contamination, and displace air bubbles (**Figure S2a**). This process was performed using 70% ethanol, phosphate-buffered saline (PBS), and 10% bleach solution. All the steps were performed at 50 μL min−1. Initially, reservoirs filled with 70% ethanol were connected to the system, and their content were flushed for 1 min. Then ethanol reservoirs were exchanged for PBS reservoirs. During each exchange, the tube was gently pressurized to expel the liquid–air interface retained at the outlet before immersion in the new reservoir. This step kept air out of the system. PBS was injected for 1 min and afterwards, the reservoirs were exchanged for 10% bleach solution reservoirs and flushed for 2 min. A second PBS wash (1 min) removed the residual bleach. Next, the system was passivated with 5% Pluronic F-127 for 2 min followed by a washing step with basal cell culture medium for 1 min. The system was then ready, and we finally exchanged each medium-filled reservoir for the corresponding working solution required for chip priming, cell pairing, and cell fusion.

Note that after the cleaning and priming of the fluidic lines, all the reservoirs are filled with cell media. Afterwards, the reservoirs are replaced with the corresponding working reagents, and the lines of the 70% ethanol, 5% Pluronic F-127 and PEG reservoirs are actuated to fill the tubing with these solutions from each reservoir to the distribution valve (**Figure S2b**). The line for the media reservoir does not require actuation at this stage because this tubing is already actuated. Also, the lines for cells are not actuated at this stage since they are used fresh and are connected only immediately before injection.

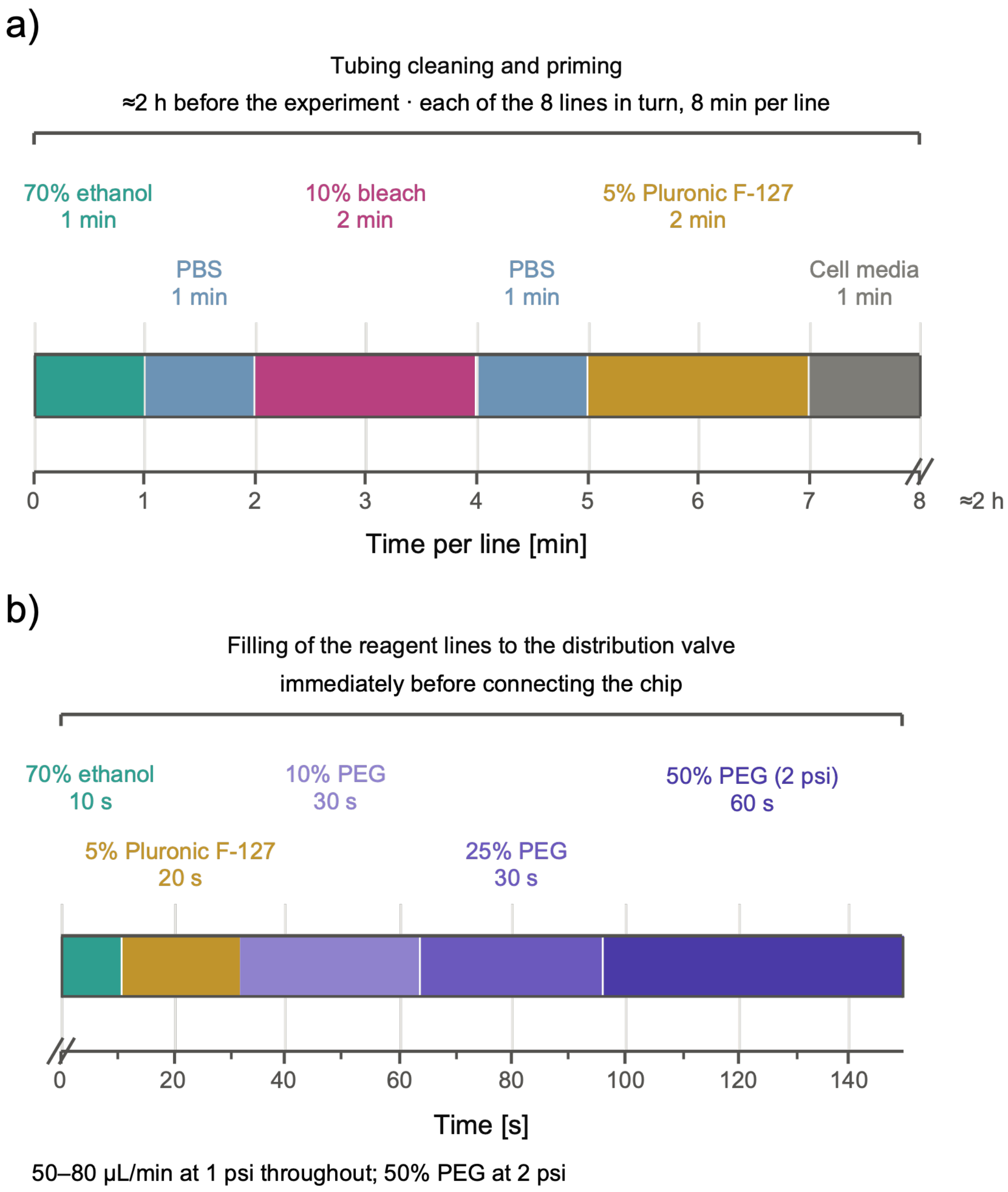


**Figure S2**. Protocol for the cleaning, priming and actuation of the tubing. a) Sequence and time steps used for the cleaning and priming of the fluidic system tubing. This was performed for each of the reagent lines. b) Filling the reagent lines with the corresponding working solution.

#### S6. Distribution of cell pairing efficiency across the circular radius of the radial chip

To corroborate if the radial chip is useful for the conditions and cell lines used here the deformability-based cel trapping and pairing, the cell pairing efficiency (trap occupancy with both cells) were calculated by a MATLAB code along the radius in **Figure S3**. It is possible to observe that the efficiency between the innermost and outermost cell trap rings is not more than 5%, which means that the variation of liquid flows along the radius is not significant for this kind of cells in the evaluated region of the cell trap area.

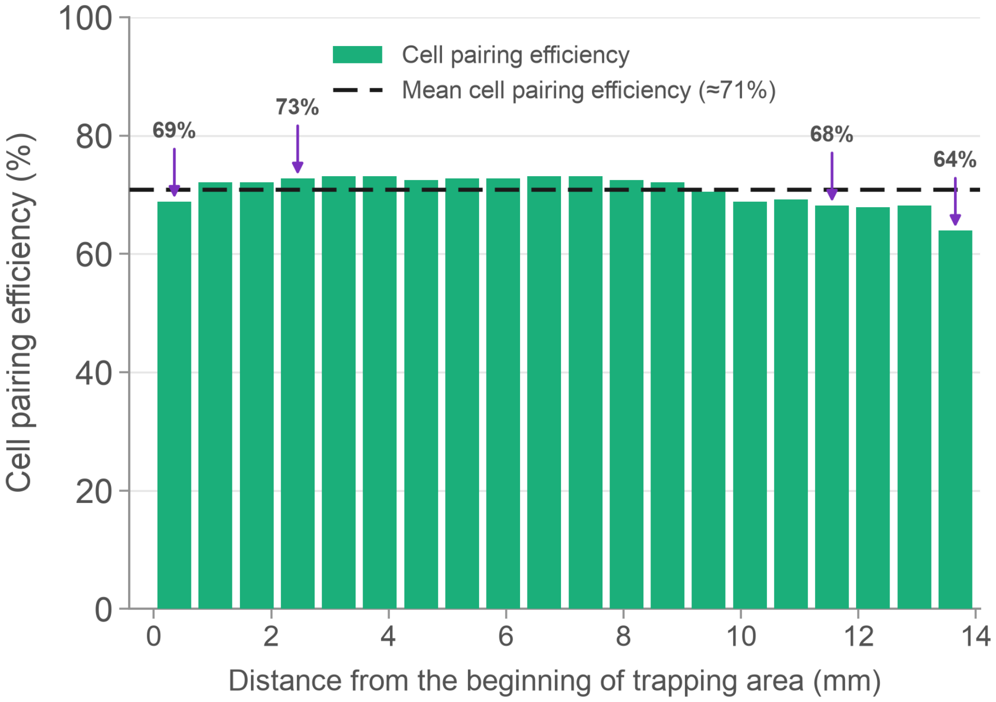


**Figure S3**. Pairing efficiency in the automated high-throughput microfluidic platform across the cell trapping rings. The percentages are calculated by a MATLAB code from the beginning of the trapping area using tiled fluorescence images.

### S7. Chemical cell fusion optimization

Flow cytometry was used to compare the step-up-and-down PEG-based protocol against their partial and single-step counterparts. All cells recovered from the chip were analyzed 24 h after fusion, and hybrids were identified as double-positive ($GFP^+/RFP^+$) events in the Q2 quadrant. **Figure S4a** shows two additional independent replicates of the step-up-and-down protocol, which yielded 18.4% and 23.4%; together with the primary run (17.7%), these give a mean of approximately 20% across three independent experiments. **Figure S4** also shows the partial and single-step comparators run on chip under otherwise identical conditions: the step-up sequence (up to 8.17%, **Figure S4b**), the step-down sequence (up to 7.10%, **Figure S4c**) and a single 50% PEG 1500 exposure (6.17%, **Figure S4d**). These measurements support the conclusion drawn in the main text that the complete step-up-and-down profile improves fusion efficiency by at least two-fold relative to any partial or single-step protocol.

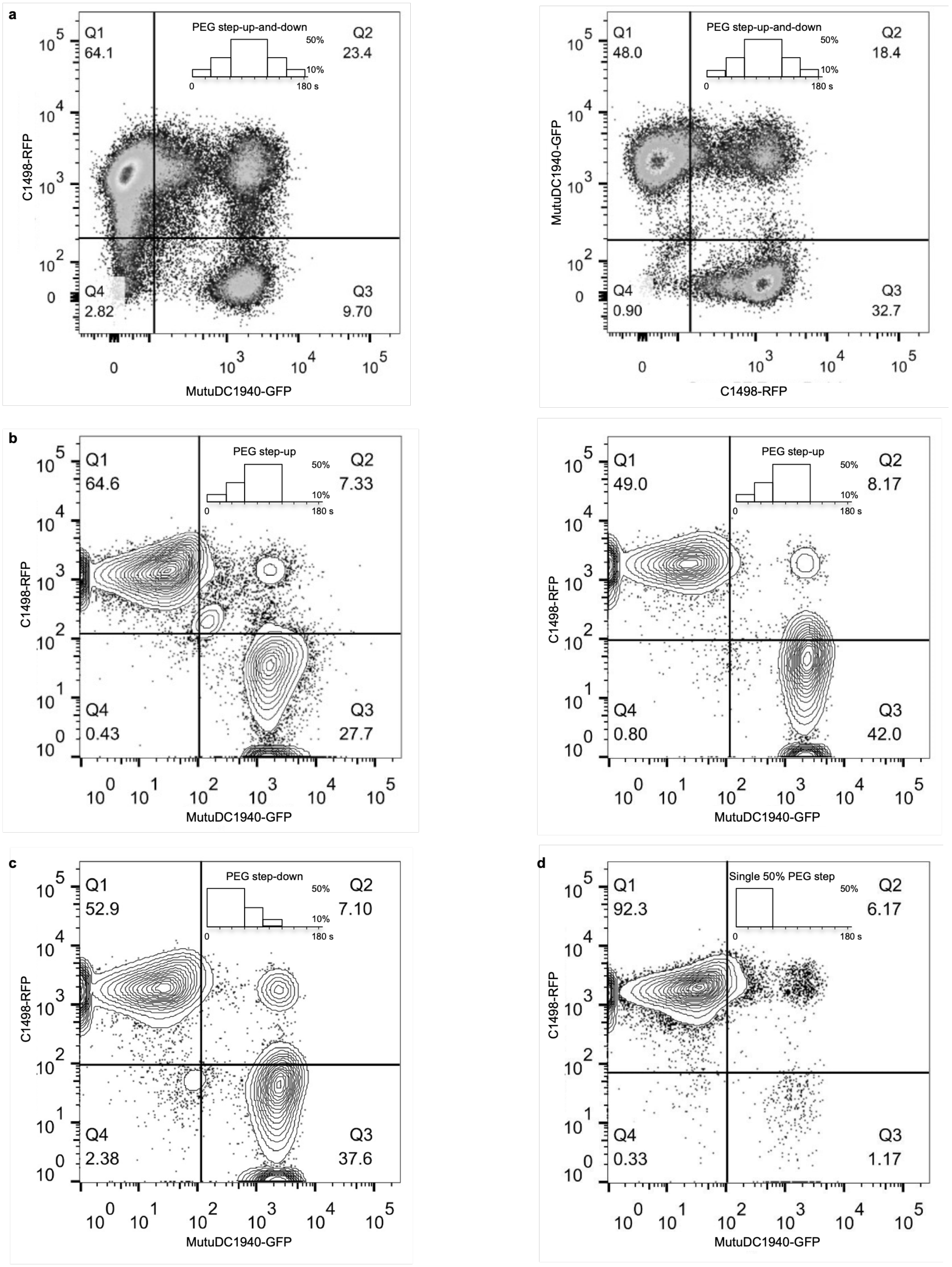

a
Q1
64.1
PEG step-up-and-down
50%
10%
0
180 s
Q2
23.4
Q4
2.82
Q3
9.70
C1498-RFP
MutuDC1940-GFP
Q1
48.0
PEG step-up-and-down
Q2
18.4
Q4
0.90
Q3
32.7
MutuDC1940-GFP
C1498-RFP
b
Q1
64.6
PEG step-up
Q2
7.33
Q4
0.43
Q3
27.7
C1498-RFP
MutuDC1940-GFP
Q1
49.0
PEG step-up
Q2
8.17
Q4
0.80
Q3
42.0
C1498-RFP
MutuDC1940-GFP
c
Q1
52.9
PEG step-down
Q2
7.10
Q4
2.38
Q3
37.6
C1498-RFP
MutuDC1940-GFP
d
Q1
92.3
Single 50% PEG step
Q2
6.17
Q4
0.33
Q3
1.17
C1498-RFP
MutuDC1940-GFP

**Figure S4**. Complementary flow cytometry results for the chemical cell fusion optimization in the high-throughput microfluidic cell pairing platform. a) Replicates of the fusion performed using the step-up-and-down PEG protocol. b) Experiments showing only the step-up PEG methodology. c) Results for the step-down PEG sequence. d) Flow cytometry results using a single 50% PEG step on chip.

### S8. Plasmid maps

Tumor cells used in the pairing and fusion experiments were fluorescently labeled so that the two partners could be distinguished within a single trap and hybrids identified by colocalization of the two channels. C1498 tumor cells were transfected to express either GFP or RFP, and the maps of the two constructs used for that transfection are provided below for reproducibility. Transfection and selection conditions are described in the Methods. **Figure S5** shows the pGFP construct carrying the GFP gene and the pRFP construct carrying the RFP gene.

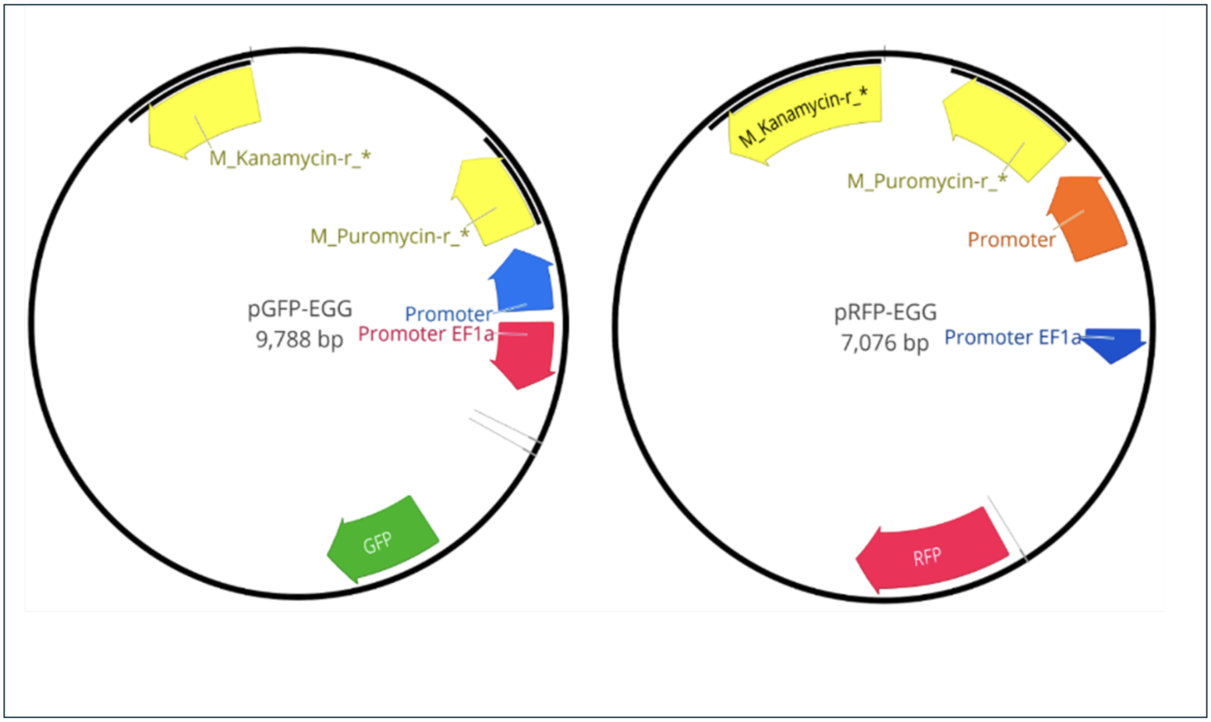


**Figure S5**. Plasmid maps used for the transfection of C1498 cells. pGFP designed plasmid containing the GFP gene and pRFP designed plasmid containing the RFP gene.
